# TorchRef: An open-source PyTorch Framework for Crystallographic Refinement

**DOI:** 10.64898/2026.05.13.724821

**Authors:** Hans-Peter Seidel, Jörg Standfuss, Tobias Weinert

## Abstract

Macromolecular crystallographic refinement underpins structural biology, yet existing software packages often lack accessible, modular codebases amenable to rapid method development. Here, we introduce TorchRef, a PyTorch-based crystallographic refinement framework that exposes all refinable parameters, atomic coordinates, displacement parameters, occupancies, and scale factors to automatic differentiation. The framework implements FFT-based structure-factor calculations, the French-Wilson treatment of intensities, bulk-solvent modeling with established mask parameters, and stereochemical restraints from the CCP4 Monomer Library. A modular target architecture allows loss functions to be combined, weighted, and extended independently of the core refinement machinery. Validation against 1,000 PDB structures demonstrates that TorchRef-based refinement reproduces a median R-free within 1% of Phenix while maintaining comparable model quality. Structure factor calculation in TorchRef scales readily across multiple CPU cores and is over 100 times faster on modern GPUs than CCTBX. To showcase how modern methods like time-resolved crystallography can benefit from the flexibility that TorchRef provides, we implemented direct refinement of a typical time-resolved model against amplitude differences, a use case currently not explored by classic refinement programs. TorchRef is released under the MIT license with full API documentation and tutorials, providing an accessible platform for developing and testing new crystallographic refinement protocols.

**Synopsis:** TorchRef is an open-source PyTorch-based crystallographic refinement framework that exposes all refinable parameters to automatic differentiation, delivers GPU-accelerated structure-factor evaluation more than 100× faster than CCTBX, and enables new workflows, such as direct refinement against amplitude differences in time-resolved crystallography.

## 1. Introduction

Macromolecular crystallographic refinement is a mature and indispensable component of structural biology and was central to building the models that inform our understanding of protein structures hosted in the Protein Data Bank (PDB)^1^. Modern refinement software such as Phenix, REFMAC5, BUSTER, and MAIN combine well-optimized model parametrization, stereochemical restraints, maximum-likelihood targets, and custom gradient calculation to strike a balance between prior knowledge and data^2–7^. These programs have been central to the success of macromolecular crystallography; however, documentation of lower-level functionality is often unavailable or difficult to access because of custom codebases that are frequently not open source.

Maximum-likelihood refinement targets, cross-validated regularization targets, and chemically informed restraint libraries stabilize refinement in challenging regimes such as low resolution, model incompleteness, and complex macromolecular assemblies. They are increasingly complemented by physically motivated force fields^3,4,6–8^. At the same time, experimental modalities have diversified. Microcrystal electron diffraction (microED) enables structure determination from nanometer-sized crystals^9,10^. X-ray free-electron lasers and upgraded synchrotrons have enabled serial crystallography (SX), which collects tens of thousands of still images from micron-sized crystals and merges them into complete datasets while mitigating radiation damage, making room-temperature measurements routinely accessible^11–13^. Time-resolved serial crystallography extends these methods to the recovery of structural intermediates spanning femtosecond-to-second timescales and has revealed reaction pathways in diverse light-triggered systems^14–17^. These modern crystallographic experiments often produce large, heterogeneous datasets that call for refinement frameworks capable of handling ensembles, mixed occupancies, and custom priors^18^.

To bridge the gap between modern crystallographic experiments, which increasingly aim to elucidate macromolecular function rather than merely determine structure, and refinement software largely built to serve broad, general-purpose needs in structural biology, a growing ecosystem of specialized tools has emerged. These include approaches that integrate molecular dynamics directly into refinement workflows^8^ or vice versa^19^, or couple quantum-mechanical calculations to crystallographic refinement^20^. Others provide methods for improving difference electron density maps^21^, which can be used for state identification^22^ or enable the refinement of weakly populated, transient states in time-resolved crystallography^23^.

In parallel, the broader machine-learning community has converged on differentiable-programming frameworks, most prominently PyTorch and JAX, that fundamentally change how numerical algorithms are developed and optimized^24–26^. In these frameworks, all numerical operations are performed on multi-dimensional arrays (tensors) and recorded in a dynamic computational graph. When a scalar objective, such as a likelihood function, is evaluated, the framework traverses the graph in reverse to compute exact gradients with respect to every parameter, a process known as automatic differentiation or backpropagation. For crystallographic refinement, this means that new target functions, parameterizations, or restraint terms can be implemented without ever deriving or coding analytical gradients by hand. Furthermore, because the same tensor operations execute on both CPUs and GPUs without code changes, algorithms that are dominated by array arithmetic, such as structure-factor summations, benefit from orders-of-magnitude acceleration on commodity GPU hardware. Within structural biology, these ideas are gaining traction^27,28^. Recent work couples machine-learning-accelerated quantum-mechanical potentials with crystallographic and cryo-EM refinement, and it has been argued that future end-to-end differentiable models should be applied in academic research^29,30^.

Building on these advances, we introduce TorchRef, a PyTorch-based macromolecular refinement framework designed primarily as a platform for method development rather than as a replacement for established refinement packages. It uses automatic differentiation to back-propagate gradients with respect to atomic coordinates, displacement parameters, occupancies, and scale factors. By combining fast Fourier transforms with PyTorch’s dynamic computational graph, the framework inherits the favorable O(N_hkl_ log N_hkl_) scaling of FFT-based crystallographic algorithms. The framework includes French-Wilson treatment of intensities^31^, bulk-solvent and anisotropic scaling following established implementation^2,4,32^, and stereochemical restraints derived from the CCP4 Monomer Library and available via the GitHub repository at github.com/MonomerLibrary/monomers. Wherever practical, functionality is implemented using PyTorch to avoid unnecessary conflicts. Full API documentation of TorchRef and tutorials are available at github.com/HatPdotS/TorchRef.

## 2. Results

### 2.1. TorchRef module structure

TorchRef implements a complete crystallographic refinement pipeline as a collection of differentiable PyTorch modules (**Fig. 1A**). The following description of the TorchRef package is not exhaustive but highlights the key design choices and features most relevant to the pipeline. For a more detailed description, we defer to the documentation available via GitHub.

**Figure 1.**
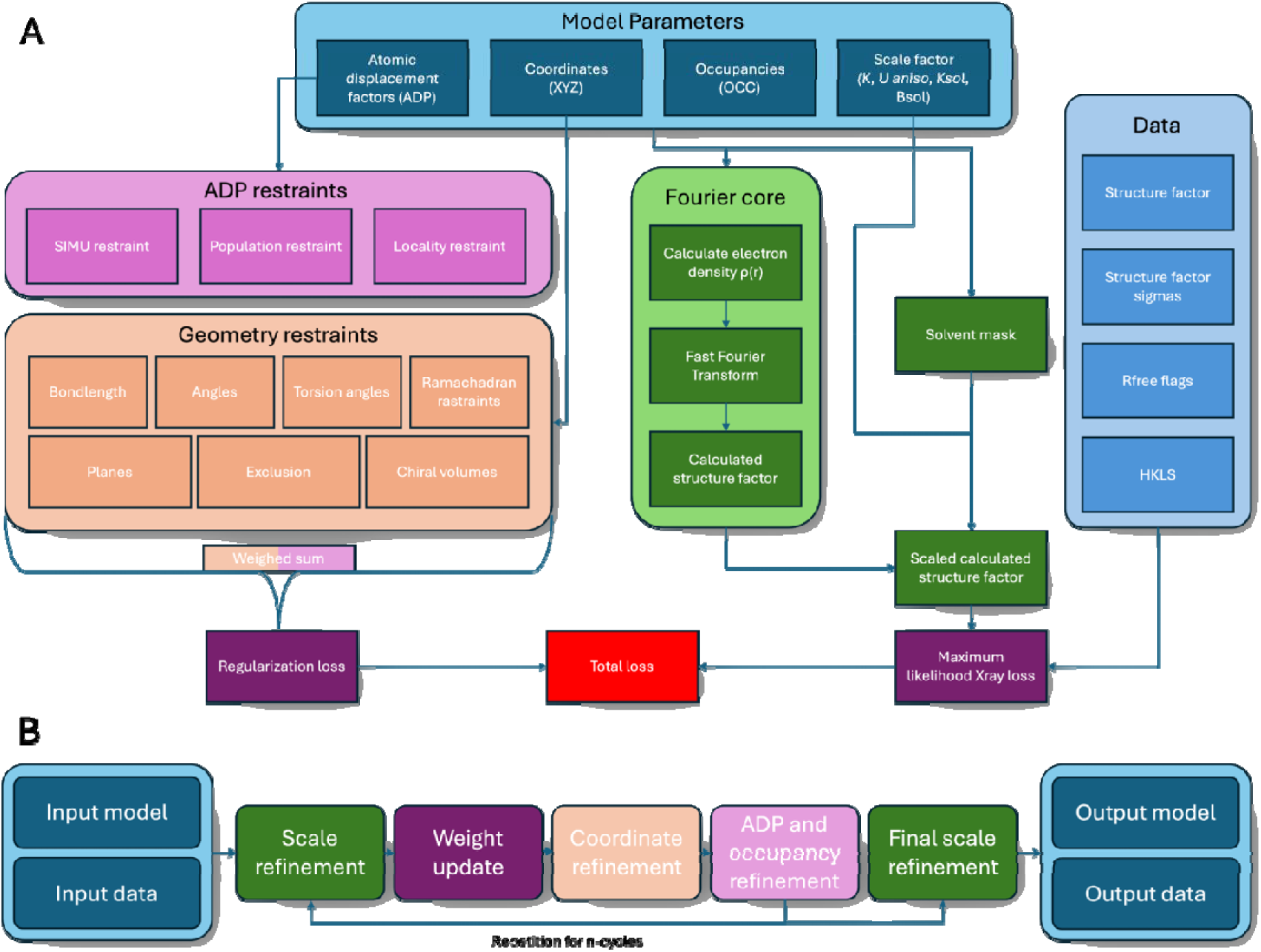
(A) Schematic overview of information flow during a forward pass in the standard TorchRef refinement loop. The central goal of any macromolecular refinement is to improve the data–model fit. Atomic coordinates are used to evaluate geometry-based restraint targets: bond lengths, bond angles, torsion angles, Ramachandran score, planarity, non-bonded repulsion, and chiral volumes. Atomic displacement parameters are used to feed three regularization targets: a SIMU-like restraint, a population-level restraint formulated as a Kullback–Leibler (KL)^39^ divergence, and a locality restraint. From all model parameters, structure factors F_calc_ are computed, scaled against experimental data, and compared with observed amplitudes F_obs_ through a maximum-likelihood target. Individual target values are aggregated according to their respective weights into a scalar loss, from which gradients are backpropagated to all refinable parameters via PyTorch’s autograd. (B) Overview of the TorchRef refinement workflow. The process begins by loading structure coordinates (PDB or mmCIF format) and reflection data (MTZ or CIF format). An initial scale refinement is performed, followed by the main refinement loop: scale update, weight update, coordinate refinement, and joint optimization of occupancies and ADPs. After the specified number of cycles, a final round of scale refinement is performed, followed by writing out the refined model and structure factors.

The base module collects all the main functionality underpinning TorchRef. Here, we collect basic input-output functions that are well optimized for both GPU and CPU. Operations are implemented wherever possible using PyTorch, enabling strong integration with its autograd framework.

Experimental data is managed by the ReflectionData data class, which tracks outliers and validation flags (R_free_ flags), handles data selection, and performs French-Wilson conversion if required. It also handles reading and writing MTZ files and writing maps in CCP4 format using the reciprocalspaceship and gemmi libraries, respectively^33,34^. Internally, we use the CCP4 convention for mapping reflections to a respective asymmetric unit. Structure factors are implemented as a masked tensor to facilitate outlier masking. We additionally implement a collection class to handle multiple datasets, including alignment and scaling.

The Model object encapsulates all refinable parameters with support for selective freezing and reparameterization. Parameter wrappers enable flexible representations: occupancies are parameterized at the residue level with optional sum-to-one constraints for alternate conformers, while ADPs are parameterized in log-space to ensure positivity. In addition to the classic parameter wrappers, TorchRef also supports internal coordinates. Here, we implement a shallow tree approach, separating trees into approximately 30 atom-rigid groups to avoid the lever-arm problem^21,35^.

The Scaler module handles bulk solvent modeling, anisotropy correction, and bin-wise scaling. Solvent masks are built using the universal parameters established by Afonine et al. (2005)^32^, with a solvent probe radius of 1.1 Å and a shrinkage radius of 0.9 Å. The scaler is fitted using PyTorch’s L-BFGS optimizer and the standard ML X-ray target. Optimization of scaling parameters is generally recommended and stable within the main refinement loop. In standard refinement, the scaler is restrained from correcting a B-factor difference between the structure and the dataset.

Geometric restraints are implemented in the Restraint module, initialized from CIF restraint files, and compute deviations from ideal geometric values. Internally, we build restraints once during initialization and cache indices required for the calculation of the respective metrics. As a CIF library, we use the CCP4 Monomer Library available on GitHub^37^.

Loss functions are implemented as dedicated target modules. The default X-ray target follows a Bhattacharyya (negative logarithmic overlap integral between two Gaussians) formulation that compares the model, accounting for its isotropic uncertainty, to the data. The model error is calculated from the position and sigma uncertainties in a Gaussian, accounting for the molecular B-factor distribution and data resolution. We also implement the classic rice-based refinement target^6^ used in other crystallographic refinement programs, a simple negative log-likelihood (NLL), and a Least Squares target. Additionally, we implement three standard ADP regularization targets, namely a locality restraint, a SIMU-like restraint (similarity restraint for ADPs of bonded atoms)^38^, and a population-level restraint, as well as seven restraint-based geometry regularization targets, namely a bond length target, an angle target, a torsion angle target, a planes target, an exclusion target, a chiral volume target, and a Ramachandran target. All regularization targets are implemented as NLLs.

The Loss state module keeps track of the respective loss functions and supports assigning weights to the different loss terms. It also implements the optimization infrastructure, handles loss pruning, automatically enables parameter gradients as needed, and integrates with automatic caching infrastructure in TorchRef.

The Refinement module (**Fig. 1B**) orchestrates the refinement macrocycles and tracks and reports refinement statistics. In agreement with Phenix, we found that the L-BFGS optimizer performs best for optimizing single-structure crystallographic models. Notably, in more complex loss-function and noisy-gradient settings, TorchRef maintains full compatibility with all other PyTorch optimizers.

### 2.2. Component weighting

Balancing the X-ray likelihood with geometric restraints and ADP regularization is a long-standing challenge in crystallographic refinement, with optimal weights that depend on data quality, resolution, and model state. As our loss functions are consequently expressed as NLLs, the most principled approach is to simply sum them. However, in practice, we observe significant overfitting as refinement advances. To combat this, we employ an overfitting weight that increases the prior weight based on the R-free gap. This keeps the worst overfitting in check and yields overfitting comparable to that of phenix-based refinement. We also investigated fine-tuning of loss components (Fig. S1) but found that while there were gains to be had, they were minor and did not justify introducing arbitrary weights. We note that the correct tuning of the strengths of different loss components is a long-standing problem in the field, and while there are empirical solutions^40^, these generally lack rational justification.

### 2.3. Validation of the refinement framework

To validate gradient flow and integration of optimization loss functions and weighting, we benchmarked TorchRef on a set of 1,000 protein structures from the PDB with resolutions ranging from 1.0 to 3.0 Å. Starting coordinates were perturbed by adding Gaussian noise with σ = 0.2 Å, and B-factors were perturbed with σ = 5.0 Å². Reflection data in CIF format were loaded using TorchRef’s I/O modules and converted to MTZ format. To ensure consistency, R_free_ flags were regenerated for all datasets. Perturbed structures were then refined against their respective datasets using both Phenix and TorchRef, with comparable protocols, yielding refined coordinates and B-factors without manual intervention. As TorchRef does not support updating water molecules, we disabled that feature in Phenix. Structures were then evaluated using REFMAC5 as an independent software package. Refinements with Phenix and TorchRef ran for ten cycles. Out of 997 starting structures (3 were missing files), TorchRef refined 978, while Phenix ran successfully for 830 structures. The main failure reasons were model interpretation issues (90), mismatches in R-free flags across Friedel pairs (40), failures in glycosidic bond determination (20), and other minor errors (17). For TorchRef with the assigned 8 GB of RAM, we encountered some out-of-memory errors on our Slurm-managed cluster (14) and other minor errors (5). Due to the low number of structures affected, these issues were not further investigated. Only structures for which refinements with Phenix and TorchRef produced results were considered for comparison.

TorchRef R_work_ and R_free_ values are, on median, 0.36 and 0.59 percentage points higher than those from Phenix refinements (**Fig. 2A**). Model geometry was assessed using bond-length and bond-angle RMSZs relative to Engh–Huber-type targets^41^ (**Fig. 2B**). Notably, TorchRef’s geometry is less tightly restrained than in Phenix by default, ADPs are more tightly regularized. Tuning geometry weights improved geometries but did not yield significant gains in R_free_ (**Fig. S1**). Comparing the structures to the pre-shaken structure, we find that TorchRef-based refinement more closely recovers the structures (**Fig. 2C**). The convergence behavior of the two refinement programs is very similar and largely determined by the structure (**Fig. S3**). Additionally, looking into resolution-dependent R-factor trends (**Fig. S2**), there is a weak trend in the R-factor difference between TorchRef and Phenix: with lower resolution, the R_free_ gap decreases, but the R_work_ gap increases, and vice versa.

**Figure 2.**
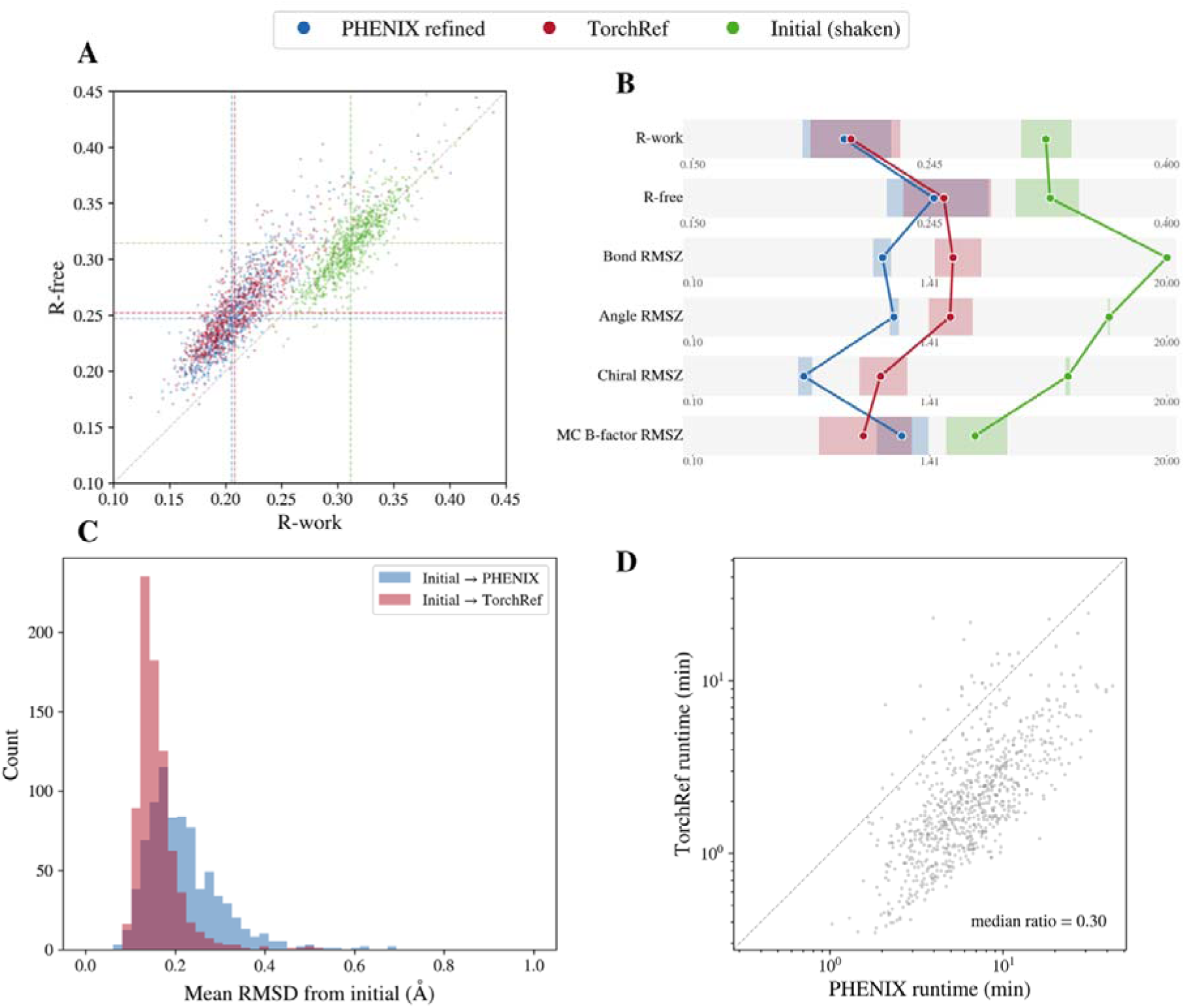
Benchmark validation on 1,000 PDB structures. Initial perturbed structures (green), Phenix-refined (blue), TorchRef-refined (red). (A) R_free_ vs R_work_ scatter plot. Dotted horizontal lines indicate median R_work_ and R_free_ values, respectively, at 0.2051, 0.2470 (Phenix), 0.2080, 0.2522 (TorchRef), and 0.3115, 0.3142 (shaken). (B) Parallel-coordinates strip chart showing various performance indices. Points indicate respective median values, while shadows indicate 25^th^ to 75^th^ percentile ranges. (C) RMSD per-structure histogram comparing the Phenix-based refinement and the TorchRef-based refinement to the original structures. (D) Scatter plot of refinement run times.

TorchRef refinements ran, on median, in 0.3× the time compared to the identical Phenix refinement on the same machine (**Fig. 2D**), likely due to PyTorch’s modern framework that scales well to multiple cores for density and gradient calculation.

### 2.4. Performance

Structure factor calculation in TorchRef is well optimized. It follows three stages. First, we splat the electron density onto the grid. This process involves calculating the per-atom electron density on a local grid and then coalescing the local grids onto a global grid. Second, we perform a fast Fourier transform on the grid. Last, we extract the structure factors from the grid in P1 and apply the symmetry operations.

The main optimization opportunity for structure-factor calculation is electron-density splatting. TorchRef’s implementation decomposes the local electron density into 1D/2D components depending on symmetry. Assuming a local electron density contribution with a radius of 5 voxels, each sphere contains 555 voxels. This approach reduces the 555×4 exponential evaluations using the ITC92 four-Gaussian approximation^42^ to 11×3×4 Gaussian evaluations for orthogonal unit cells. For monoclinic and hexagonal, we calculate 2D correction factors for each plane using 11×11×4 Gaussian evaluations per additional plane. In the triclinic case or for anisotropic scattering, we fall back to the trivial splatting approach. This optimization significantly reduces computational load in orthogonal cells and still provides measurable uplift in monoclinic and hexagonal cells. On the GPU, we use a custom Triton kernel^43^ that implements this approach; however, the performance uplift over the trivial approach is negligible (**Fig. S4**). Numerically, gradients between the trivial and the decomposed approach differ, as the trivial approach splats spheres compared to encompassing cubes in the decomposed case. In real-world cases, gradients match far below the noise level.

With this optimization in place, the CPU performance of the structure factor calculation on a single CPU core is comparable to the structure factor calculation with CCTBX (**Fig. 3A**). TorchRef’s structure factor calculation scales with diminishing returns across multiple CPU cores. Based on this, TorchRef by default configures four cores in normal environments (automatically detecting available cores on Slurm). At four cores, TorchRef’s implementation is >2× faster than CCTBX. When running on the GPU with the custom Triton kernel, structure factor calculation is >100× faster in TorchRef compared to CCTBX (**Fig. 3A**).

**Figure 3.**
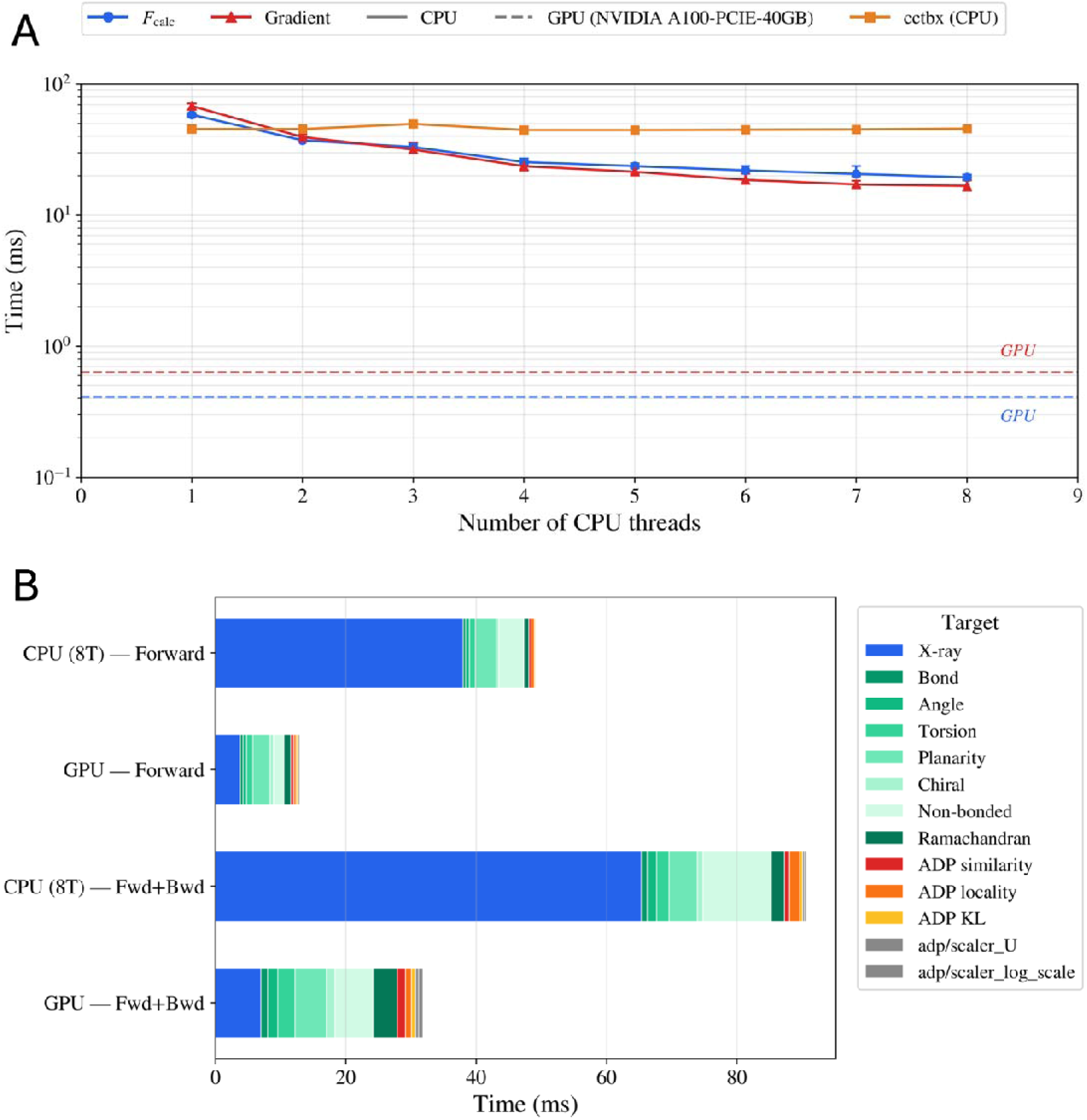
Various performance benchmarks for a 3,051-atom model (1DAW, 23,356 reflections, d_min_ = 2.05 Å). GPU timings were recorded on an NVIDIA A100 40GB. (A) Wall-clock time per F_calc_ evaluation as a function of CPU thread count. The TorchRef forward pass (blue) and backward pass (red) are shown alongside the CCTBX structure factor calculation (orange). The gradient calculation shows identical scaling behavior to the structure factor calculation and is slightly but measurably faster on the CPU. Dashed horizontal lines indicate GPU timings. TorchRef surpasses CCTBX on 2 CPU threads and scales further with more CPU cores. TorchRef achieves a >100× speedup on the GPU compared to CCTBX. (B) Comparison of loss function evaluation times for both the forward (loss calculation) and the backward pass (gradient calculation). A forward and backward pass on the GPU takes ∼32 ms and ∼92 ms on the CPU when using 8 cores.

When evaluating real-world loss functions, current implementations incur significant overhead. A full refinement cycle takes ∼92 ms on 8 CPU cores and ∼32 ms on the GPU (**Fig. 3B**). While this is far from the potential ∼1 ms cycle time of the F_calc_ calculation, it still results in sub-second L-BFGS steps (assuming 10 evaluations per step). Due to the diverse nature of the refinement targets and the existing data-loading overhead, we did not optimize the target implementations. Notably, when many F_calc_ values need to be calculated, as during ensemble refinement, we do not need to compute loss functions for each molecule; instead, we can aggregate structure factors prior to loss evaluation.

### 2.5. Difference Refinement

The end-to-end differentiability of TorchRef enables gradient propagation through complex computational pipelines that would be difficult to implement in conventional refinement programs. This explicitly allows for multi-model multi-dataset workflows. As an example, we implement difference-amplitude-based refinement, which is useful for time-resolved crystallography and related workflows. In addition to the regularization loss functions, geometric restraints, and atomic-displacement-based regularization, we add an X-ray difference loss based on a negative log-likelihood formulation. The difference target explicitly explains differences between two related datasets, thereby avoiding systematic errors that affect both datasets. Additionally, we add a similarity loss that penalizes the light-to-dark model for small changes, assuming that small changes are likely noise, but large changes are real. In practice, this requires a well-refined dark model, as errors in the dark model will propagate into the difference.

Combining this with mixed-model refinement, where we define a combined model as a linear combination of a reference and a light model, we can optimize the light model to explain the observed structure factor differences. This is a simple example of a multi-model, multi-dataset approach that can be extended to fit the data. The modular structure of TorchRef and its inherited automatic backpropagation enable more complex scenarios in which a set of mixed models can be refined that combine a small number of structures across many datasets, for instance, a time series of a reaction with changing intermediate populations.

To validate different solutions, we implemented a command-line tool that calculates local correlation coefficients between the weighted difference observed structure factor (WDF_o_) density map and the weighted difference calculated structure factor (WDF_c_) density map for a given atom selection and masking radius (see Materials and Methods) under consideration of the dark phases. Here, we use inverse-variance weights for the difference observations. Additionally, we calculate reciprocal-space correlation on the differences. Localization of the signal within the structure results in significantly higher real-space correlations than reciprocal-space correlations, enabling meaningful correlations to be extracted even for a strongly localized change.

To assess the utility of the difference-refinement approach, we applied it to dataset 7YYZ from a time-resolved serial crystallography study of the photo-pharmacological drug azo-combretastatin A4 (azo-CA4) bound to αβ-tubulin^44^ (for the full refinement protocol, see Materials and Methods). In that study, cis-to-trans photoisomerization of the azobenzene moiety was triggered by light, and nine structural snapshots spanning 1 ns to 100 ms captured the resulting ligand release and collapse of the colchicine-site binding pocket. The 7YYZ dataset corresponds to the 10 µs timepoint, at which the ligand has begun to shift but remains bound. The deposited structure was originally refined in Phenix using extrapolated structure factors at an estimated light-state occupancy of 0.22.

TorchRef difference refinement was performed with a resolution cutoff of 2.2 Å at an assumed light state occupation of 0.22, matching the published structure. The resulting model matched the WDF_o_ with a local correlation of 0.847, compared to 0.735 for the deposited model refined using extrapolated structure factors (**Fig. 4**) within a 2.5 Å radius around both the dark and light conformations of the IBL ligand (azo-CA4). Considering the global fit of the model, the calculated difference structure factor amplitudes show correlations of 0.509 and 0.310 with the observed difference structure factor amplitudes for the TorchRef difference-refined model and the extrapolated model, respectively. Notably, the differences in the resulting structures are subtle and the models are qualitatively similar.

**Figure 4.**
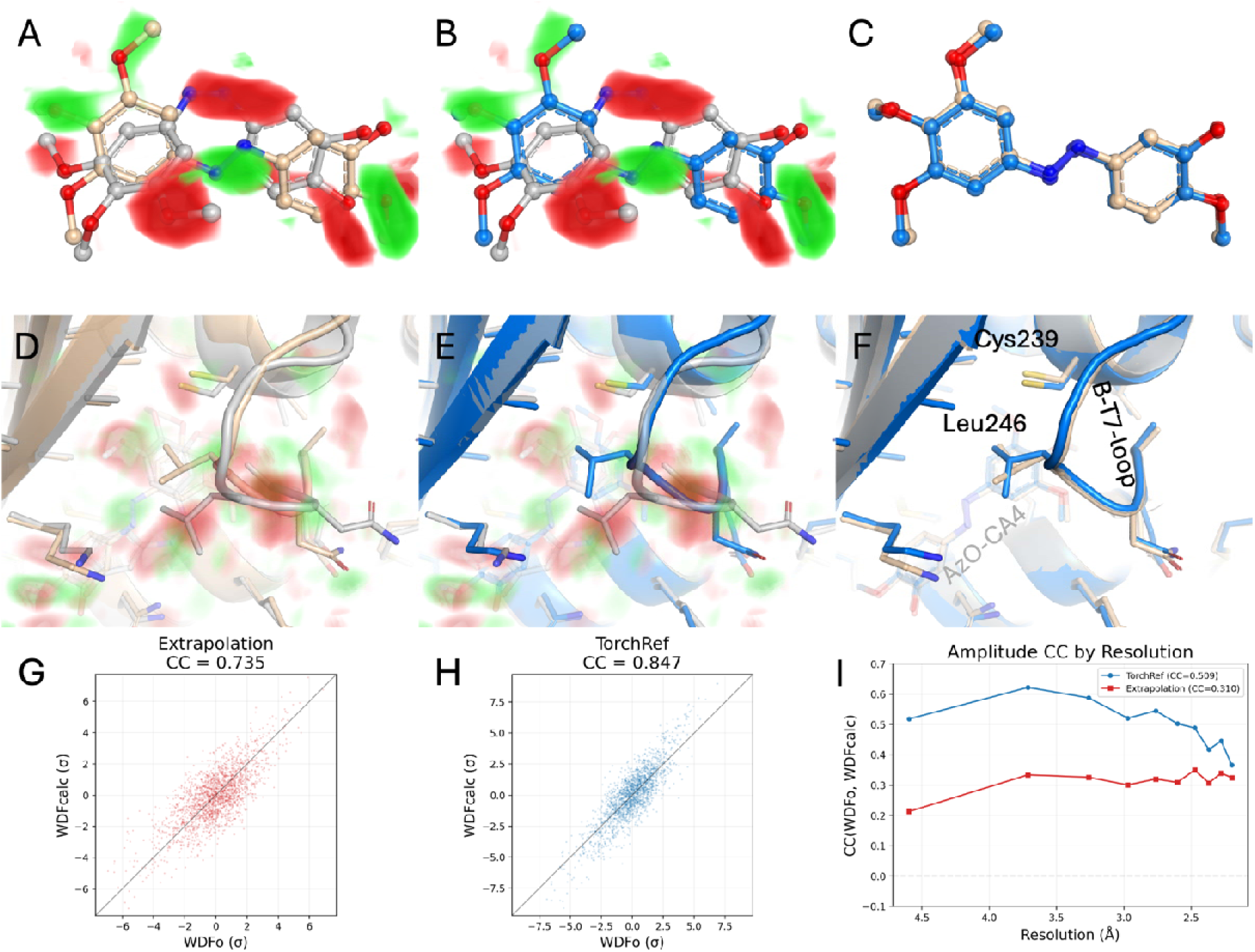
Difference refinement of azo-CA4 (IBL) *cis*-to-*trans* isomerization in tubulin (7YYZ). Weighted difference electron density (WDF_o_) maps are visualized via a volume ramp from 3 to 5 sigma around the ligand (green, positive; red, negative) (A, B) and 2.5 to 4.5 sigma in the wider binding pocket (D, E). Top row (A–C): IBL ligand close-up. Bottom row (D–F): binding pocket view showing the changes in the βt7-loop resulting from the unbinding of the *trans* conformer ligand. Dark-state structure 8QL2 (gray), published extrapolated structure 7YYZ (wheat), and TorchRef difference-refined structure (blue). (A, D) WDF_o_ map with the deposited 7YYZ structure refined from extrapolated structure factors. (B, E) WDF_o_ map with the TorchRef difference-refined structure. (C, F) Superposition of the TorchRef-refined (blue) and deposited 7YYZ (wheat) conformations, showing the agreement and differences between the independently determined light-state structures. Scatter plots G and H show the distribution of WDF_c_ voxel values relative to WDF_o_ voxels within a 2.5 Å radius around the dark and light conformations of the ligand. G shows the scatter plot for the extrapolated data, while H shows the TorchRef difference-refinement scatter plot. The correlation between calculated and observed difference amplitudes is shown in panel I, binned by resolution.

## 3. Discussion

TorchRef provides a PyTorch-based crystallographic refinement framework that exposes all relevant parameters to automatic differentiation. This design paradigm offers significant advantages over classic implementations, as it correctly handles complex gradient computations through automatic backpropagation, removing the need for manually derived analytical gradients. This not only reduces implementation effort but also eliminates an important source of programming errors when developing new refinement targets or parameterizations. TorchRef’s native PyTorch implementation enables deep integration with the growing ecosystem of PyTorch-based tools in structural biology, including molecular dynamics engines such as TorchMD, structural prediction models such as OpenFold, and machine-learning-accelerated quantum-mechanical potentials such as those used in AQuaRef^30^.

Our benchmark against 1,000 PDB structures demonstrates that TorchRef produces models with R-factors within 1 percentage point of Phenix’s median, while maintaining comparable stereochemical quality. Given that the two programs differ in outlier rejection, scaling implementation, the restraint library used, and the considerable optimization effort invested in Phenix over many years, this modest offset is expected and falls well within the range commonly observed between established refinement packages. Median refinement times were 0.3× those of Phenix, likely reflecting the efficient parallelization of modern tensor frameworks.

The difference-refinement procedure demonstrates a use case that is difficult to implement in conventional refinement programs. Existing approaches to time-resolved crystallographic data analysis typically rely on extrapolated structure factors, which amplify the signal from a partially occupied excited state to simulate a fully occupied one. While powerful, extrapolation introduces noise amplification that scales inversely with occupancy.

Additionally, because extrapolated structure-factor amplitudes are calculated with phases derived from the dark state, any phase difference between the dark and light models introduces error into the resulting maps. Furthermore, extrapolation systematically produces negative amplitudes (due to phase mismatches) that require correction, potentially introducing bias. In practice, this error behaves predominantly as additional noise rather than as a systematic bias. This explains the general success of extrapolation-based methods but also means that the excited-state density in the active region is systematically noisier than the true signal. Furthermore, to obtain more agreeable statistics, modern extrapolation tools apply Bayesian weighting to downweigh noisy-difference amplitudes, which, while reducing noise, introduces additional dark-state bias by shrinking the signal toward the reference^23^.

Direct refinement of a mixed model against amplitude differences, as implemented in TorchRef, sidesteps these issues by fitting the difference between the dark-state model and the occupancy-weighted combination of dark- and light-state models to the observed difference in the dark and light data. In the case of 7YYZ, this approach yielded a higher local correlation between calculated and observed difference maps (0.847 vs. 0.735 for the deposited structure around the chromophore).

Notably, the differences between the extrapolated structure and the TorchRef-difference refined structure are subtle; however, quantitatively, this represents a significant increase in agreement between the model and the data. Additionally, we believe this approach has potential for extension in the future, including consideration of more than two datasets and multiple transient structures.

TorchRef is designed as a platform for method development and does not yet implement the full feature set of established refinement packages. At present, it lacks automated water placement and updating, NCS restraints, twinning support, and TLS parameterization. Simulated annealing, which can help escape local minima during early refinement stages, is not implemented, though TorchRef’s compatibility with any PyTorch optimizer means stochastic gradient-based alternatives could serve a similar role.

Given the well-documented, open-source nature of TorchRef, we hope to make the development of custom refinement protocols accessible to a broader community of crystallographers and method developers. The full API documentation, Jupyter notebook tutorials, and permissive MIT license are intended to lower the barrier to experimentation. As a concrete example, the difference refinement was initially implemented in approximately 1000 lines of Python code (core logic takes up 200 lines), a task that would require substantially more effort in a conventional refinement codebase, where analytical gradients would need to be derived and implemented manually.

Looking ahead, TorchRef’s modular architecture opens several avenues for development. The multi-model, multi-dataset framework demonstrated here for difference refinement generalizes naturally to kinetic multi-state refinement against time-resolved serial crystallography data, where a small number of conformational states would be refined against many datasets simultaneously. Integration with neural network potentials is a particularly compelling direction: replacing empirical geometric restraints with learned energy surfaces such as MACE-OFF or AIMNet2 would combine physically motivated priors with the data-driven X-ray maximum likelihood, much as AQuaRef^30^ has demonstrated for quantum-mechanical refinement within Phenix, but with the advantage that, in TorchRef, the potential is a native PyTorch module and its gradients propagate through the same computational graph identical to other targets of the refinement.

More broadly, TorchRef’s GPU-accelerated structure factor calculation, which is two orders of magnitude faster than conventional implementations, opens use cases where rapid structure evaluation is needed, including training deep learning models on observed structure factors, ensemble refinement with full gradient updates, and on-the-fly map computation during manual model building.

We hope that the open-source, modular, and well-documented nature of TorchRef will allow many more researchers to contribute effectively to macromolecular structure refinement, a field in which the complexity of established codebases and the absence of accessible documentation have historically concentrated methodological development among a small number of individuals. We believe this will help address long-standing challenges, such as integrating physical theory with experimental data and the persistent gap between model and observation, reflected in R-factor statistics that have remained qualitatively unchanged despite decades of methodological refinement.

## 4. Materials and methods

### 4.1. Structure selection and dataset preparation

A non-redundant benchmark dataset of 1,000 protein crystal structures was assembled from the Protein Data Bank (PDB). Structures were selected programmatically via the RCSB PDB Search API using the following criteria: (i) experimental method restricted to X-ray diffraction, (ii) resolution between 1.5 and 3.0 Å, and (iii) maximum pairwise sequence identity of 30%, enforced by retaining only cluster representatives from the RCSB sequence clustering at the 30% identity level. From an initial pool of 1,500 candidate entries, structures were filtered to retain only those with deposited structure factor data, yielding the final set of 1,000 structures.

For each structure, coordinate files (PDB and mmCIF formats) and structure factor files (SF-CIF format) were downloaded from the RCSB. Ligand restraint dictionaries were obtained from the RCSB Chemical Component Dictionary for all non-standard residues, excluding common solvents and ions (HOH, WAT, NA, CL, K, CA, MG, ZN, FE, SO4, PO4, GOL, EDO).

### 4.2. MTZ conversion and R-free flag generation

Structure factor CIF files were converted to MTZ format using TorchRef’s ReflectionData module. To ensure unbiased cross-validation, R_free_ flags were regenerated for all structures (regenerate_rfree_flags(force=True)), establishing a consistent and independent test set partition across all entries. This is necessary because deposited R_free_ flags carry memory of the original refinement and cannot be used for unbiased comparison between different refinement programs.

### 4.3. Generation of perturbed starting models

Perturbed models were generated by deliberately perturbing each deposited structure (“shaken”) to create degraded starting models. Coordinate shifts were sampled from a Gaussian distribution with a sigma of 0.2 Å, and isotropic B-factors were perturbed by shifts randomly sampled from a Gaussian distribution with a sigma of 5.0 Å², using TorchRef’s Model.shake_coords and Model.shake_b_factors methods, respectively. This ensures that both refinement programs start from identical, suboptimal models, preventing either from benefiting from proximity to the deposited coordinates.

### 4.4. Reference refinement with PHENIX

Phenix refinement was performed using PHENIX version 1.20-4459^45^. Each shaken structure was refined against the corresponding MTZ file containing the regenerated R_free_ flags. The refinement protocol consisted of 10 macrocycles with the following settings: refinement strategy individual_sites + individual_adp + occupancies (simultaneous refinement of atomic coordinates, individual isotropic B-factors, and partial occupancies); simulated annealing disabled; target weight optimization disabled (both XYZ and ADP weight optimization set to false); bulk solvent and scaling enabled; ordered solvent disabled; Ramachandran restraints enabled. For structures containing non-standard residues, geometry restraints were either sourced from pre-computed restraint libraries or generated de novo using phenix.elbow with geometry optimization (--opt --do-all). Malformed CRYST1 records (containing “None” as Z-value) were automatically corrected before refinement. Jobs were submitted to a Slurm cluster with four CPU cores and 8 GB of RAM per structure.

### 4.5. TorchRef refinement

TorchRef refinement was performed using the torchref.refine command-line script. Default weights were used. Jobs were submitted using four CPU cores and 8 GB of RAM per job. Refinement was run on TorchRef v0.5.0.

### 4.6. Validation

Validation metrics were computed independently of both refinement programs using third-party crystallographic tools. R_work_ and R_free_ values were calculated using REFMAC5^3^ from the CCP4 suite (version 8.0) with zero refinement cycles (NCYCLES 0, MAKE HYDR NO). This provides an independent, unbiased assessment of model-to-data agreement without further modifying the refined coordinates.

Stereochemical quality was assessed from the REFMAC5 zero-cycle output, which reports RMS deviations from ideal geometry for bond distances, bond angles, chiral volumes, planar groups, and non-bonded contacts (van der Waals and hydrogen bonds). ADP geometry metrics, including main-chain B-value RMSDs, were also extracted.

Deviations between deposited models, TorchRef-refined models, and Phenix-refined structures were quantified using per-atom RMSD values after matching atoms by PDB identifier and aggregated into per-structure mean RMSD values. We collected reported wall times from log files for successful refinement runs.

### 4.7. Benchmarking structure factor calculation and gradient performance

#### 4.7.1. Test system

All benchmarks used the crystal structure of 1DAW (PDB entry 1DAW), comprising 3,051 atoms and 23,356 unique reflections at a resolution of 2.05 Å.

#### 4.7.2. Quantities benchmarked

Three timing scenarios were measured for TorchRef, each calling compute_structure_factors directly (bypassing the model cache): (1) Forward: Structure factor calculation F_calc(h) inside a torch.no_grad context, measuring the raw forward-pass cost without constructing a computational graph. (2) Backward only: A forward pass followed by loss.backward() (where loss = sum of |F_calc|), with only the backward pass timed; this isolates the cost of automatic differentiation through the structure factor kernel. This timing is less accurate than combined forward+backward pass evaluations due to timing overhead. (3) Forward + backward: End-to-end F_calc computation plus gradient back-propagation, timed together. As a reference, the forward-only F_calc computation via CCTBX (xray_structure.structure_factors(d_min=…).f_calc()) was timed on CPU under identical conditions. We did not collect gradients for CCTBX because the documentation was not clear on how to do so.

#### 4.7.3. CPU thread scaling

To assess parallelism, benchmarks were run across 1–8 CPU threads. Each configuration was executed in an isolated subprocess with TORCHREF_NUM_THREADS, OMP_NUM_THREADS, MKL_NUM_THREADS, and OPENBLAS_NUM_THREADS all set to the target thread count, ensuring consistent control over intra-op parallelism in PyTorch, BLAS, and OpenMP. The TorchRef_init module reads TORCHREF_NUM_THREADS and calls torch.set_num_threads() accordingly.

#### 4.7.4. GPU benchmarks

GPU timings were collected on an NVIDIA A100-PCIE-40GB. All GPU timings bracket each iteration with torch.cuda.synchronize() to ensure asynchronous kernel launches are fully completed before reading the wall-clock time.

#### 4.7.5. Timing protocol

For each configuration (thread count × device), three warm-up iterations were run (discarded) followed by 10 timed iterations. Wall-clock time was measured with time.perf_counter(). Summary statistics reported are the mean, minimum, and maximum over the 10 timed iterations. Speedup is defined as the ratio of the single-thread CPU mean time to the mean time at a given thread count (or on GPU).

#### 4.7.6. Execution environment

Each thread-count configuration was run in a separate Python subprocess to avoid cross-contamination of the thread pool state. The orchestrator script (benchmark_thread_scaling.py) launches worker processes sequentially via subprocess.run, collects per-run JSON results, and aggregates them into a summary CSV.

#### 4.7.7. Model setup

The ModelFT class was initialized with a standard radius_angstrom=3.0 (real-space summation cutoff) at the experimental resolution. Isotropic and anisotropic parameters (xyz, ADP, occupancy, scattering factors) were extracted via M.get_iso() / M.get_aniso(), cloned, and detached with requires_grad=True to ensure a fresh autograd graph for each backward-pass iteration.

#### 4.7.8. Loss function timing

Loss function timings were measured using TorchRef’s standard loss function initialization as done in a standard refinement setup. We then evaluated the forward and backward passes for all registered targets using the LossState.targets dictionary.

### 4.8. Difference refinement protocol

Difference Refinement was performed using the torchref.difference-refine command-line script. In the simplest case implemented here, the dark model is considered frozen, which aids refinement stability and reduces computation time. The optimization targets comprise the classic regularization targets for geometry and atomic displacement parameters, the X-ray ML target for the mixed model against the light data, and the difference target. Optimization proceeds according to an annealing schedule that progressively reduces the weight of the difference target (default: 5, 3, 2). This is done to encourage the model to escape the local minima of the optimized geometry in the dark model. The light-state fraction is not refined by default, as it is correlated with the atomic positions of the light structure; fraction optimization can be enabled via a command-line flag. Refined models and corresponding structure factors are written out to facilitate validation and manual correction in model-building software such as Coot (for a full description of outputs, see the documentation).

For validating difference-refined models, we use the torchref.validate-ded command-line script, which calculates difference-amplitude correlation metrics between the weighted F_obs_ data and the structure factor differences computed from the mixed and dark models. The primary metric is a local correlation coefficient between the weighted difference observed structure factor (WDF_o_) map and the weighted difference calculated structure factor (WDF_c_) map, computed within a specified masking radius around a given atom selection. Maps were calculated using dark-state phases.

To demonstrate the difference-refinement approach, we refined against pdb entry 7YYZ^44^, a time-resolved serial crystallography structure of a photoisomerized azobenzene analog ligand. The deposited structure was originally refined in Phenix using extrapolated structure factors with an occupancy of 0.22 at a resolution cutoff of 2.2 Å. For TorchRef difference refinement, we used the recently released dark structure 8QL2 as both the initial light and dark models, with a fixed starting occupancy of 22%. Following an initial round of automated refinement, sections of the model were manually adjusted in real space using Coot, using the phase-corrected, extrapolated structure factors written out by torchref.difference-refine, to accelerate convergence. Two cycles of alternating manual model building and torchref.difference-refine were performed. In a final refinement run the xray weight was reduced to 1 to equilibrate the structure into a favorable geometry. The refinement used reflections to a resolution cutoff of 2.2 Å. Validation correlations were computed at 2.2 Å resolution to allow direct comparison with the deposited model. R-free flags were grafted from the 8QL2 dataset to the 7YYZ dataset to allow for meaningful R-free values.

## Acknowledgements

The project was conceived by H.-P.S. and T.W. The code was authored by H.-P.S. under the supervision of T.W. and J.S. The manuscript was authored by H.-P.S. and T.W. with input from J.S. Figures were made by H.-P.S. with the help of T.W. The manuscript was reviewed by J.S., T.W. and H.-P.S. Claude Code was extensively used during the development of TorchRef for prototyping, refactoring, and documentation. Claude was used to review and rephrase text during the preparation of this manuscript. This work was supported by Swiss National Science Foundation project grants 10.007.643 (J.S.) and 10.003.847 (T.W.).

## Data availability

TorchRef is available on GitHub under an MIT license and can be installed via pip. Full API documentation and Jupyter notebook tutorials are available via GitHub at github.com/HatPdotS/TorchRef. The scripts used to generate the figures are available via GitHub. The difference-refined structure is pending acceptance in the PDB and will be available as under code 30AS.

## Supporting information

Supplementary figures (**Figs. S1–S4**) provide additional validation data supporting the main-text results.

**Figure S1.**
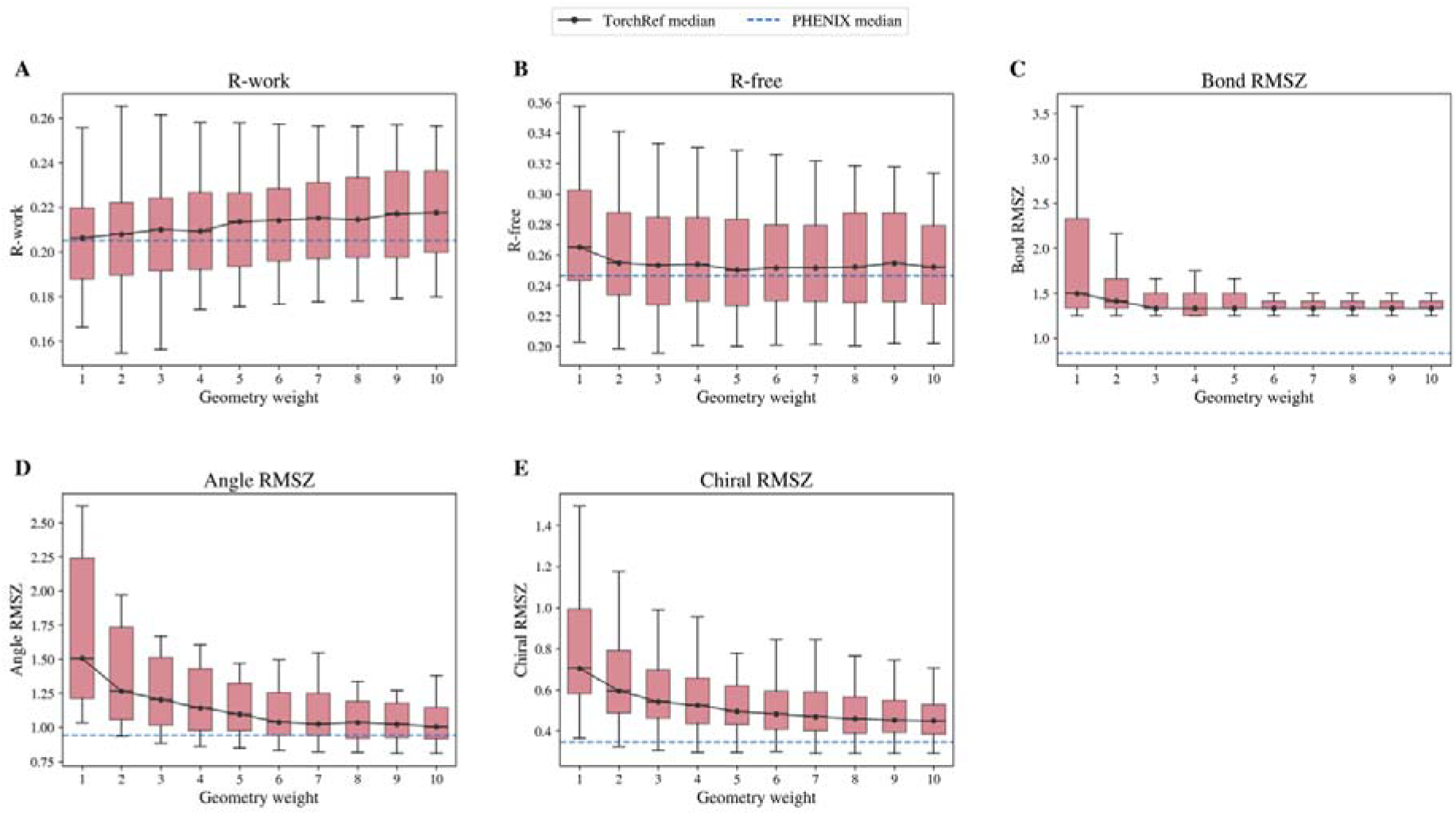
Box-plot panel showing refinement quality as a function of the geometry restraint weight multiplier (1–10) shown for a random subset of 50 structures. Five panels display the distributions of R-work, R-free, Bond RMSZ, Angle RMSZ, and Chiral RMSZ across 50 benchmark structures. PHENIX medians from the main validation benchmark are overlaid as dashed reference lines.

**Figure S2.**
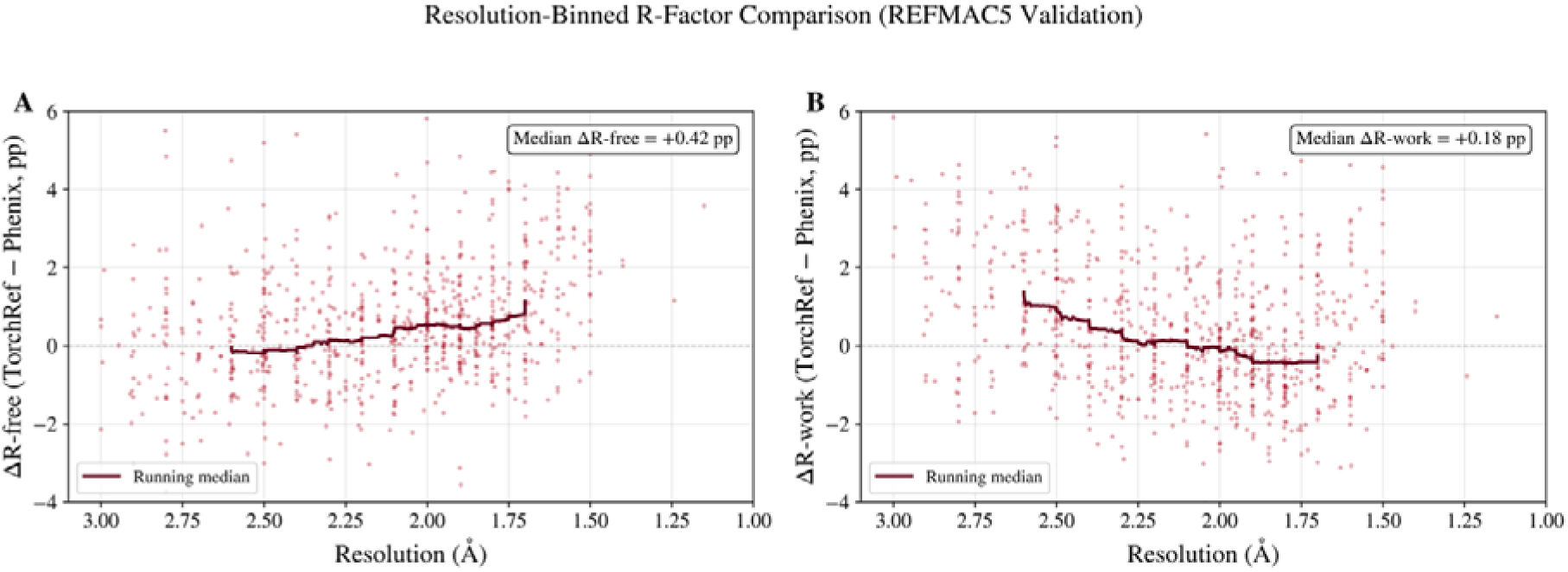
Two-panel scatter plot showing the per-structure R-factor gap (TorchRef minus PHENIX) as a function of resolution. Panel A: ΔR-free vs resolution. Panel B: ΔR-work vs resolution. Running medians are overlaid.

**Figure S3.**
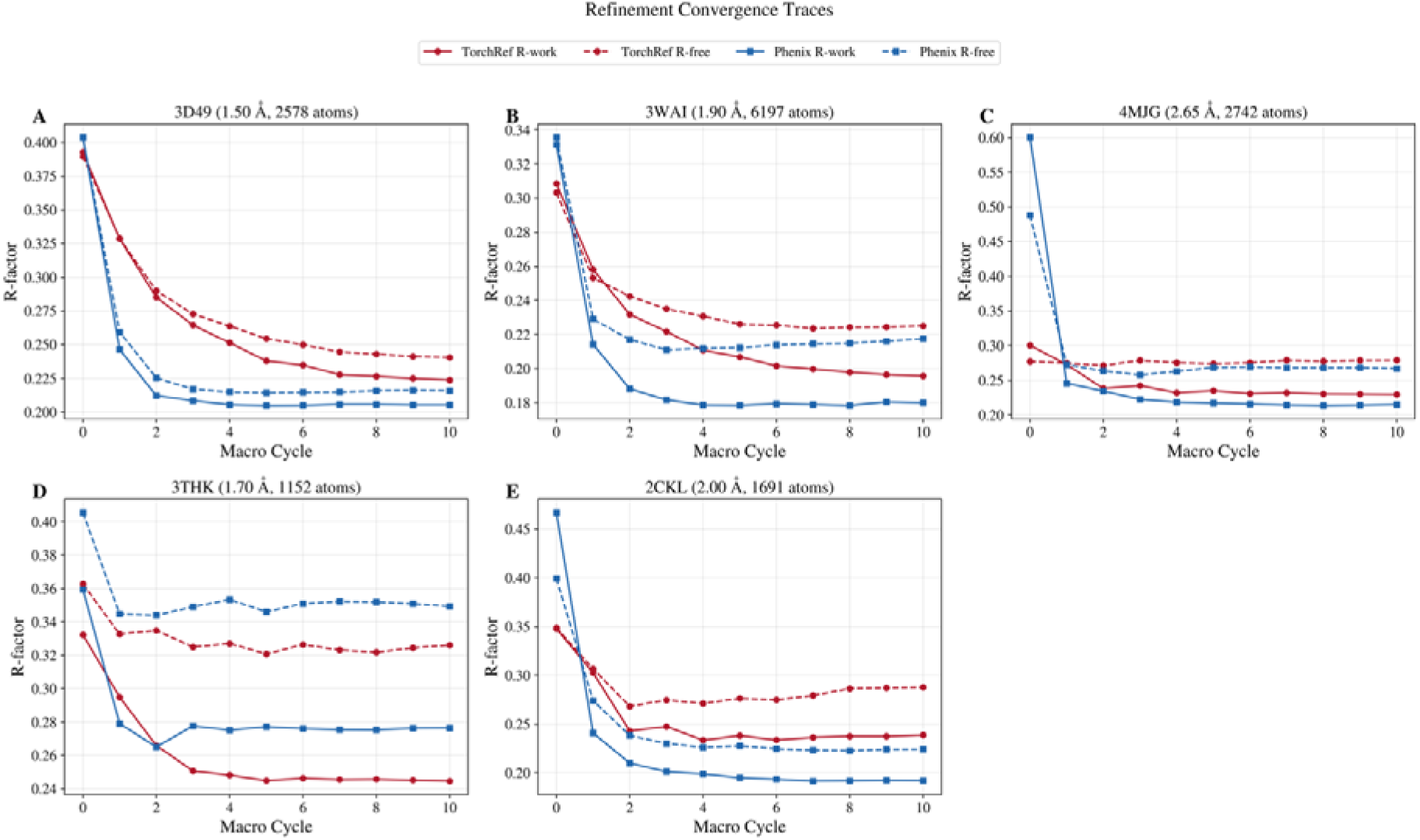
Grid of panels (one per representative structure) showing per-macrocycle R-work and R-free for both TorchRef and PHENIX, starting from the same shaken model. Structures span the resolution range (high, medium, low) and include cases where each program outperforms the other. Demonstrates smooth, monotonic convergence and comparable convergence rates.

**Figure S4.**
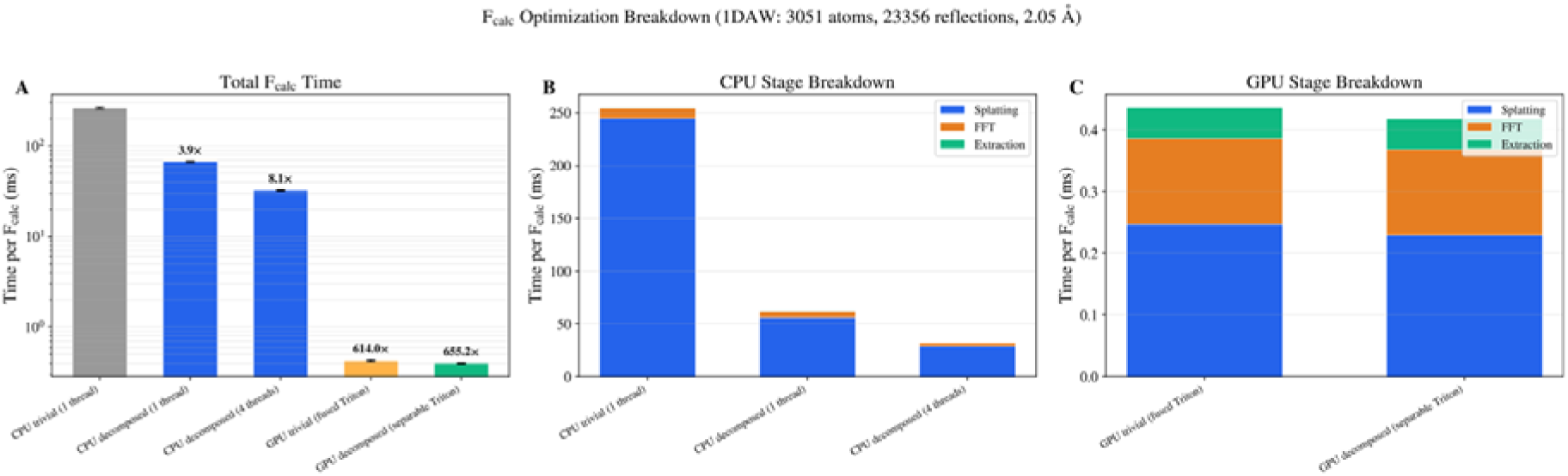
Three-panel figure comparing five F_calc configurations on the 1DAW benchmark structure. Panel A: total F_calc wall time per approach on a log scale (CPU trivial, CPU decomposed 1-thread, CPU decomposed 4-thread, GPU fused Triton, GPU separable Triton) with speedup annotations. Panel B: CPU stage breakdown (splatting, FFT, extraction) as stacked bars. Panel C: GPU stage breakdown. Quantifies the contribution of the decomposed splatting approach and the Triton GPU kernel.

